# Durga: An R package for effect size estimation and visualisation

**DOI:** 10.1101/2023.02.06.526960

**Authors:** Md Kawsar Khan, Donald James McLean

## Abstract

1. Statistical analysis and data visualisation are integral parts of science communication. One of the major issues in current data analysis practice is an over-dependency on – and misuse of – *p*-values. Researchers have been advocating for the estimation and reporting of effect sizes for quantitative research to enhance the clarity and effectiveness of data analysis. Reporting effect sizes in scientific publications has until now been mainly limited to numeric tables, even though effect size plotting is a more effective means of communicating results, however, statistical software for plotting effect sizes is currently limited.
2. We have developed the Durga R package to estimate and plot effect sizes. Durga allows users to estimate unstandardised and standardised effect sizes and bootstrapped confidence intervals of the effect sizes. The central functionality of Durga is to combine effect size visualisations with traditional plotting methods.
3. Durga is a powerful statistical and data visualisation package that is easy to use, providing the flexibility to estimate effect sizes of paired and unpaired data using different statistical methods. Durga provides a plethora of options for plotting effect size, which allows users to plot data in the most informative and aesthetic way.
4. Here, we introduce the package and its various functions. We further describe a workflow for estimating and plotting effect sizes using example data sets.

## Introduction

Null hypothesis significance testing (NHST), despite being extensively criticised by researchers, has long been the most popular statistical approach for data analysis (Coe, 2002; Stunt et al., 2021). NHST tests a null hypothesis against an alternative hypothesis to reject or accept the hypothesis based on a *p*-value (Dushoff et al., 2019; Nickerson, 2000). Yet, *p*-values can be misleading and have several limitations. *P*-values cannot be directly compared between studies and often trigger unjustifiable false comparisons (Bernardi et al., 2016; Berner & Amrhein, 2022; Halsey, 2019). A statistical significance indicated by a *p*-value of less than 0.05 is often erroneously misinterpreted as indicating a meaningful effect size, whereas statistically non-significant results often have an underlying non-zero effect size (Bernardi et al., 2016; Berner & Amrhein, 2022; Halsey, 2019). Furthermore, use of *p*-values and NHST effectively asks the binary question, “is there an effect?”, whereas studies in ecology and evolution are typically quantitative and “how large is the effect?” is usually a more important question (Ho et al., 2019; Sullivan & Feinn, 2012). Therefore, researchers are advised to move away from a *p*-value imposed binary decision making to quantitative analyses, and the use of estimation statistics is rightfully gaining momentum (Amrhein et al., 2019; Muff et al., 2022). Many statistical packages such as SPSS, MATLAB, Python and R now enable researchers to estimate effect sizes.

In addition to recommending the use of estimation statistics over NHST, researchers and statisticians have also been advocating plotting effect sizes alongside traditional plots (Cumming, 2012; Gardner & Altman, 1986). Conventional plots depicting group data as bar charts, box plots or violin plots, may include the group mean ± error bar, and can indicate statistically significant differences between groups with an asterisk (“*”). Although conventional chart types can provide information about group data distribution, range, central tendency and deviation from central tendency, they do not convey the main information of interest, i.e., the differences between groups — the effect size. Researchers therefore suggest plotting effect size along with group data, and replacing the statistical significance indicator “*” with confidence intervals of effect size. Gardner–Altman (1986) suggested plotting effect size to the right of the group plot and Cummings (2012) suggested that multiple effect sizes could be shown beneath the group plot. Estimation graphics communicate quantitative differences between groups i.e., effect size, thereby making data interpretation much easier. Despite being such a powerful tool for data visualisation, good software and applications for making varied and aesthetic estimation plots are lacking. Existing estimation plotting software, including the R package ‘dabestr’ (Ho et al., 2019), the statistical software GraphPad and the ESCI package (https://thenewstatistics.com/itns/esci/), provide some plotting options, but producing plots beyond the capabilities of these packages is complex and time consuming.

Here, we describe Durga, an R package for effect size estimation and visualisation. Using Durga, researchers can easily estimate and plot unstandardised or standardised effect sizes for paired and unpaired data. Standardised effect sizes can be calculated using Cohen’s d and Hedges’ g. Durga also estimates bootstrapped confidence intervals of effect sizes. Durga is the most powerful graphical tool for plotting effect sizes available, and provides researchers with a simple yet flexible means to plot effect size within the same graphic as traditional comparative graphs such as box plots, violin plots, bar plots and mean-error plots.

### Durga

Durga defines two primary functions, `DurgaDiff()` and `DurgaPlot()`. Durga is written within base R, consists of entirely new code, and provides users with a large range of options for plotting group data (bars, boxes, violins, central tendency, error bars, individual data points, and all possible combinations of them) and effect sizes together with their confidence intervals (below or to the right of the group data, display or hide bootstrap violins, control display symbology); alternatively, effect size confidence intervals can be plotted above group data in the form of confidence brackets. Durga is fully compatible with the multiple plot layout mechanisms of base R; `par(mfrow = c(…))`, `layout()` and `split.screen()`. By defining sensible defaults for most options, users of the package can explore the options they are interested in, while ignoring functionality that is not currently relevant. While the existing R package `dabestr` builds on `ggplot` to provide effect size estimation and plotting (Ho et al., 2019), group data display is limited to grouped scatter plots. Durga aims to provide greater plotting flexibility and creative power with an interface that is easy for non-expert R users to understand and use, eliminating the need to master `ggplot`. Durga plots are highly modifiable, which provides a flexible and creative interference to plot informative as well as aesthetic plots.

Durga is implemented in R (R Core Team, 2022). The current version of the package (1.0) requires R ≥ 4.2.0 and is submitted in CRAN and currently under review; soon could be installed from CRAN (https://cran.r-project.org/web/packages/Durga/index.html) and directly via the R console using *install*.*packages(‘Durga’)*. The development version of the package is available for download through GitHub (https://github.com/KhanKawsar/EstimationPlot) and can be installed using `devtools::install_github(“KhanKawsar/EstimationPlot”, build_vignettes = TRUE)`. The package has been developed using R packages boot (Canty & Ripley, 2021), RColorBrewer (Neuwirth, 2022) and vipor (Sherrill-Mix & Clarke, 2017).

### DurgaDiff ()

The `DurgaDiff` function estimates between-group effect sizes. Data to be analysed must be in a data frame (or similar) organised in *long format*, which means there is a row for each observation, a column for the measured value datum and another that identifies the observation treatment or group. The `data.col` argument to `DurgaDiff` specifies the data column and `group.col` specifies the group column. Group values need not be numeric. Columns may be identified by name or index. Effect sizes can be calculated for either paired or unpaired data. When the data are paired, the `id.col` argument is required to specify the identity of each individual datum or specimen.

`DurgaDiff` calculates standardised or unstandardised effect sizes; the desired effect type is specified with the argument `effect.type`. Unstandardised effect size is calculated as the mean difference between compared groups and is specified with `effect.type “mean”`. Standardised effect sizes can be calculated as Cohen’s d or Hedges’ g, and are specified with `effect.type “cohens”` or `“hedges”` respectively. DurgaDiff uses equation 1 (below) to calculate Cohen’s d and equation 2 to calculate Hedges’ g for unpaired data, where *M*_*i*_, *SD*_*i*_ and *n*_*i*_ are the mean, standard deviation and sample size of group *i*. Equation 3 (below) is used to calculate Cohen’s d (d_z_) for paired data (Lakens, 2013), where the numerator, *M*_*diff*_, is the mean of the difference scores, the denominator is the standard deviation of the difference scores (*X*_*diff*_) and is the number of scores.

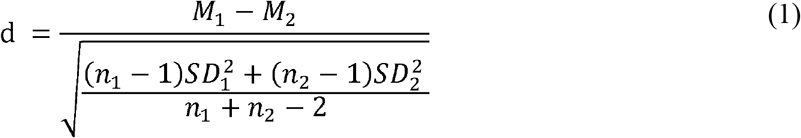

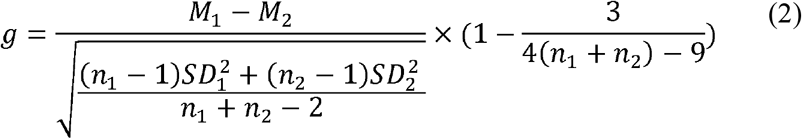

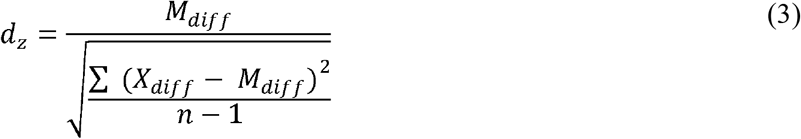

Confidence intervals for the estimate are determined using bootstrap resampling, using the adjusted bootstrap percentile (BCa) method, calculated using the `boot` and `boot.ci` functions from the boot package (Canty & Ripley, 2021). The number of bootstrap replicates may be specified by the argument `R`, and confidence level by the argument `ci.conf`.

The default behaviour of `DurgaDiff` is to order the groups alphabetically, then calculate differences between all pairs of groups. The order of groups and the labels used to represent groups can be altered by the argument `groups`, while `contrasts` can be specified to change the pairs to be compared and/or the direction of comparisons. A detailed description of the `DurgaDiff` function, including detailed descriptions of `contrasts` and `effect.type`, is available via the `DurgaDiff` help page (run the R command “`?DurgaDiff`” to view it).

The function DurgaDiff returns an object of class `DurgaDiff`, which is a list containing multiple named elements. Full details are available on the help page, however two important elements are “`group.statistics`” and `“group.differences”`. Element `group.statistics` is a matrix that summarises the groups, with a row for each group and columns for sample mean, median, standard deviation (sd), standard error (se), confidence interval (CI.lower and CI.upper) and sample size (n). “`group.differences`” is a list of `DurgaGroupDiff` objects, each of which contains bootstrapped confidence interval information for one contrast. An object returned from DurgaDiff can be used for effect size plotting using the function DurgaPlot.

#### Box1

input and output of the DurgaDiff function

~~~
library(Durga) ## Load data
data(“damselfly”)
DurgaDiff(damselfly, data.col = 1, group.col = 3,
effect.type = “cohens”, na.rm = TRUE)
## Output
Bootstrapped effect size
length ∼ group
Groups:
mean median sd se CI.lower CI.upper n
adult 32.26985 32.354 0.9919583 0.1462563 31.97527 32.56442 46
juvenile 31.16274 31.196 0.8240126 0.1479970 30.86049 31.46499 31
Unpaired Cohen’s d (R = 1000, bootstrap CI method = bca):
juvenile - adult: -1.19245, 95% CI (bca) [-1.72593, -0.731584]
~~~

### DurgaPlot()

The `DurgaPlot` function plots group data and estimated effect size, based on the result of a previous call to `DurgaDiff`. The effect size is displayed as a point – the sample statistic – together with a vertical bar representing the confidence interval of the statistic. The effect size is displayed using a different y-axis scale than the group data, which is depicted by a secondary y-axis. The y origin represents zero difference between groups, so if the confidence interval does not overlap 0, there is a significant difference between groups. Additionally, the bootstrapped distribution of the statistic is drawn as a violin plot (which is truncated at the extents of the confidence interval by default). Each effect size represents the difference between a pair of groups. Effect size display is controlled by the argument *ef*.*size*; if `FALSE`, effects sizes will not be displayed. A single effect size (i.e., when only two groups have been compared) can be plotted on the right side as suggested by Gardner-Altman (1986), in which case the secondary y-axis is shown on the right of the plot (Figure 1). Multiple effect sizes may be shown below the group data, as suggested by Cumming (2012), with the secondary y-axis shown to the left of the effect sizes (Figure 3). The position of the effect size is specified by the argument `effect.size.position`, which must be either `“right”` or `“below”`. Display of the bootstrapped effect size violin plot can be controlled by the `ef.size.violin` argument. The `contrasts` argument can be used to select which effect sizes are to be plotted; this is particularly useful for multiple groups where user might not wish to plot all possible pairwise effect sizes.

**Figure 1:**
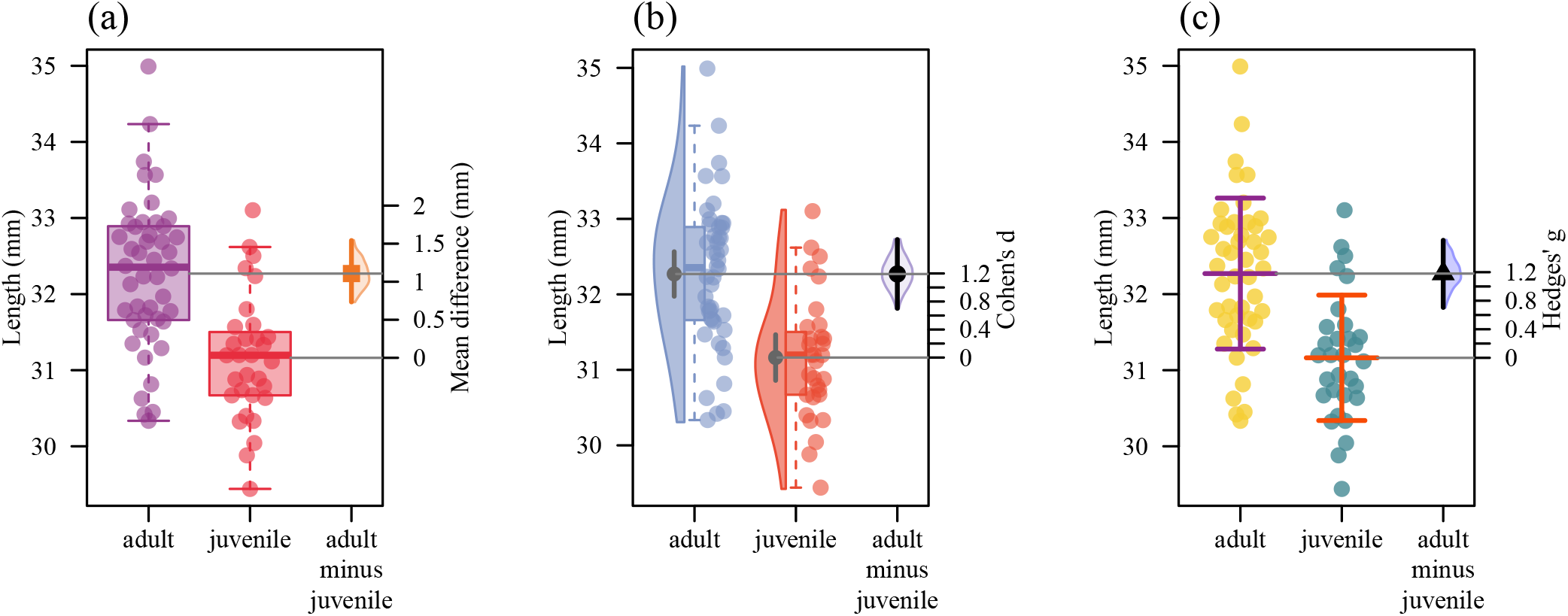
Gardner-Altman plot showing differences in body size between adult and juvenile damselflies. Left axis represents group data and right axis represents effect size. Horizontal lines are drawn from the means of each group. A) Left two boxplots depict group data, the half violin on the right exhibits the distribution of bootstrapped differences, the solid square shows mean difference, while the vertical bar shows 95% confidence interval of mean difference. The boxplots display the group median and the 75th and 25th percentiles. The whiskers extend to the minimum and maximum, but exclude outliers that are beyond 1.5 times the interquartile range. Circles on the boxes indicate individual values. B) Left two boxplots and half violins exhibit group data and the violin on the right exhibits the distribution of bootstrapped Cohen’s d (standardised mean difference), black circle represents Cohen’s d, and vertical bar shows 95% confidence interval of Cohen’s d. Half violins on the left represent the group data distribution. Grey circles inside violins represent group means, and the vertical bars through the circles represent 95% confidence intervals of group means. The boxplots display the group median and the 75th and 25th percentiles. The whiskers extend to the minimum and maximum values, but exclude outliers that are beyond 1.5 times the interquartile range. Circles adjacent to boxes indicate individual values. C) Left horizontal and vertical bars and points exhibit group data, and the half violin on the right shows the distribution of bootstrapped Hedges’ g (standardised mean difference), triangle shows Hedges’ g and vertical bars show 95% confidence interval of Hedges’ g. Horizontal line within the group data represents mean, and vertical line represents standard deviation. Circles indicate individual values.

The `DurgaPlot` function provides users with a range of options to visualise group data: box plots (argument `box=TRUE`), bar charts (`bar=TRUE`) and violin plots (`violin=TRUE`). Individual data points can be plotted (`points=TRUE`) and visually arranged using different algorithms (argument `points.method`). Control over display of the mean or median of each group is provided by the argument `central.tendency`, and control over display of the confidence interval of the central tendency by the argument `error.bars`.

`DurgaPlot` differs from other data visualisation packages in the flexibility and versatility it provides, coupled with its ease of use. Group plot representations such as box plots, bar charts and violin plots can be selected or omitted by simply by specifying `TRUE` or `FALSE` for the appropriate arguments. Additionally, multiple plot types, for example box plots and violin plots, can be combined into a single plot (Figure 1, 3). Furthermore, central tendency and error bars can be over-plotted on the combined plot (Figure 1, 3). Finally, positions of the bar, box, violin and central tendency can be shifted along the x-axis using `bar.dx`, `box.dx`, `violin.dx` and `central.tendency.dx`. This flexibility and diversity of options makes `DurgaPlot` very powerful, and provides the opportunity to produce a wide range of plots that are currently used for data visualisation across different research fields (see Supporting Information S1 for sample figures, and Supporting Information S2 for code to produce the figures). `DurgaPlot` can also be used to make traditional plots without plotting effect size (`ef.size=FALSE`) (see Supporting Information S3 and S4 for sample figures, and Supporting Information S2 for code to produce the figures). Additional details on the DurgaPlot function, including detailed descriptions of each argument and how to use them, are available on the DurgaPlot help page. The package vignette demonstrates some of the many possible plots and how to produce them, with R code included.

### Other functions

Durga provides two further functions: `DurgaTransparent()` and `DurgaBrackets()`. `DurgaTransparent` is a utility function that adds (or removes) transparency to a colour. For example `DurgaTransparent(“red”, 0.75)` returns the colour red with 75% transparency.

`DurgaBrackets` annotates an existing Durga plot with confidence brackets. Confidence brackets depict confidence intervals between pairs of groups by visually joining them with a horizontal bar and displaying the confidence interval as text. Confidence brackets portray less information than full effect sizes, but may be appropriate when many effect sizes need to be shown on a plot. Refer to the `DurgaBrackets` help page and the package vignette for more details and examples of use.

### Example usage

#### Example 1: calculate and plot two sample unpaired and paired data

Here we demonstrate the functionality of `DurgaDiff`, `DurgaPlot`, `DurgaBrackets` and `DurgaTransparent` on previously published data. For unpaired data we used the body length of juvenile and adult male damselflies (installed as the data set “damselfly” with the Durga package) (Khan & Herberstein, 2021). We first calculated effect size of the difference between the two groups using the `DurgaDiff` function. Researchers might calculate unstandardised (mean difference), Cohen’s d or Hedges’ g effect types for this analysis; we calculated all three types to demonstrate how Durga can be used to calculate different effect sizes. We then plotted the three different effect sizes together with group data using the `DurgaPlot` function (Figure 1a-c; R code in the Supporting Information S2).

To report the result, researchers may examine the `DurgaDiff` output and write that: adult damselflies (n = 46, M = 32.26, SD = 0.99) have larger body sizes than juvenile damselflies (n = 31, M = 31.16, SD = 0.82) (mean difference: 1.10, 95% CI [0.71, 1.54], Figure 1a), or (Cohen’s d = 1.19, 95% CI [0.66, 1.60], Figure 1b), or (Hedges’ g = 1.18, 95% CI [0.64, 1.69], Figure 1c).

#### Example 2: calculate and plot two sample paired data

For paired data we used the blood glucose levels of rabbits before and after administering insulin (available as the data set “insulin”) (Banting et al., 1922). We calculated unstandardised (mean difference) and standardised (Cohen’s d and Hedges’ g) effect sizes for the paired data and produced three different plots (Figure 2a-c; R code in the Supporting Information S2).

**Figure 2:**
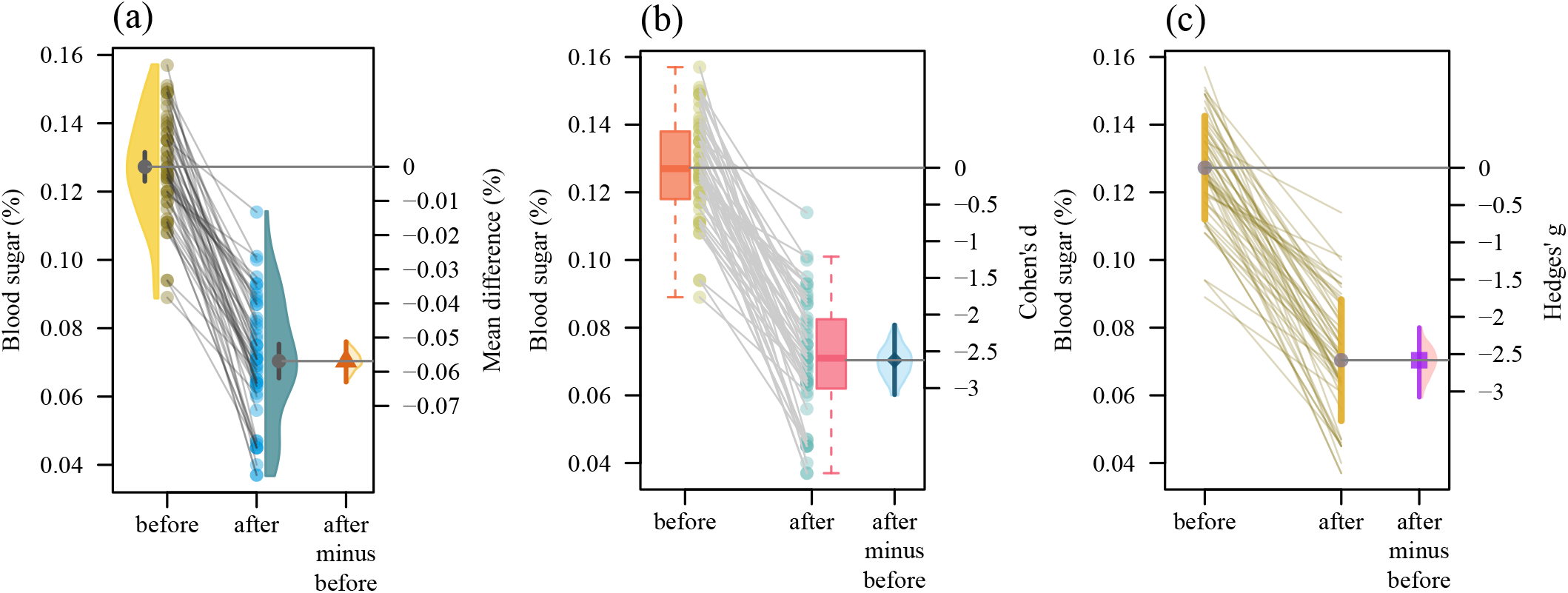
Gardner-Altman plot showing difference of blood glucose level before and after administering insulin. Left axis represents group data and right axis represents effect size. Horizontal lines extending to the right axis are drawn from the means of each group. A) Left half violins represent group data distributions and right half violin exhibits effect size statistics. Circles inside violins represent group means, and vertical bars through the circles represent 95% confidence intervals of the group means. Circles adjacent to violins indicate individual measurements, grey lines connect measurements of each individual. Half violin on the right represents the distribution of bootstrapped differences, the solid triangle shows mean difference and vertical bar shows 95% confidence interval of mean difference. B) Left two boxplots exhibit group data, and right violin exhibits effect size statistics. The boxplots display the group median and the 75th and 25th percentiles. The whiskers extend to the minimum and maximum values, but exclude outliers that are beyond 1.5 times the interquartile range. Circles adjacent to boxes indicate individual values. Grey lines connect measurements of each individual. Violin on the right exhibits the distribution of bootstrapped Cohen’s d (standardised mean difference), circle represents Cohen’s d, and vertical bar shows 95% confidence interval of Cohen’s d. C) The two circles indicate group means and the vertical bars through the circles represent 95% confidence intervals of the group means. Light brown lines connect measurements of each individual. Half violin on the right shows the distribution of bootstrapped Hedges’ g (standardised mean difference), square shows Hedges’ g, and vertical bars through the square show 95% confidence interval of Hedges’ g.

Researchers could describe the results as: blood glucose level was measured in 52 rabbits, which showed that blood glucose was lower after insulin administration (M = 0.070, SE = 0.002) than before administration (M = 0.13, SE = 0.002) (mean difference: 0.06, 95% CI [0.05, 0.06], Figure 2a); or (Cohen’s d = 2.62, 95% CI [2.13, 3.09], Figure 2b); or (Hedges’ g = 2.58, 95% CI [2.06, 3.01], Figure 2c).

#### Example 3: Calculate and plot group data with more than two groups

We further used Durga to calculate effect sizes and visualise three groups using Charles Darwin’s plant height measurements of self-fertilised, cross-fertilised and Westerham-crossed plants (available as the data set “petunia”) (Darwin, 1900). First, we used `DurgaDiff` to calculate pairwise differences between the three plant cross types. We then applied `DurgaPlot` to visualise the pairwise differences. Figure 3 shows three possible ways to visualise the group differences and effect sizes (R code in the Supporting Information S2).

**Figure 3:**
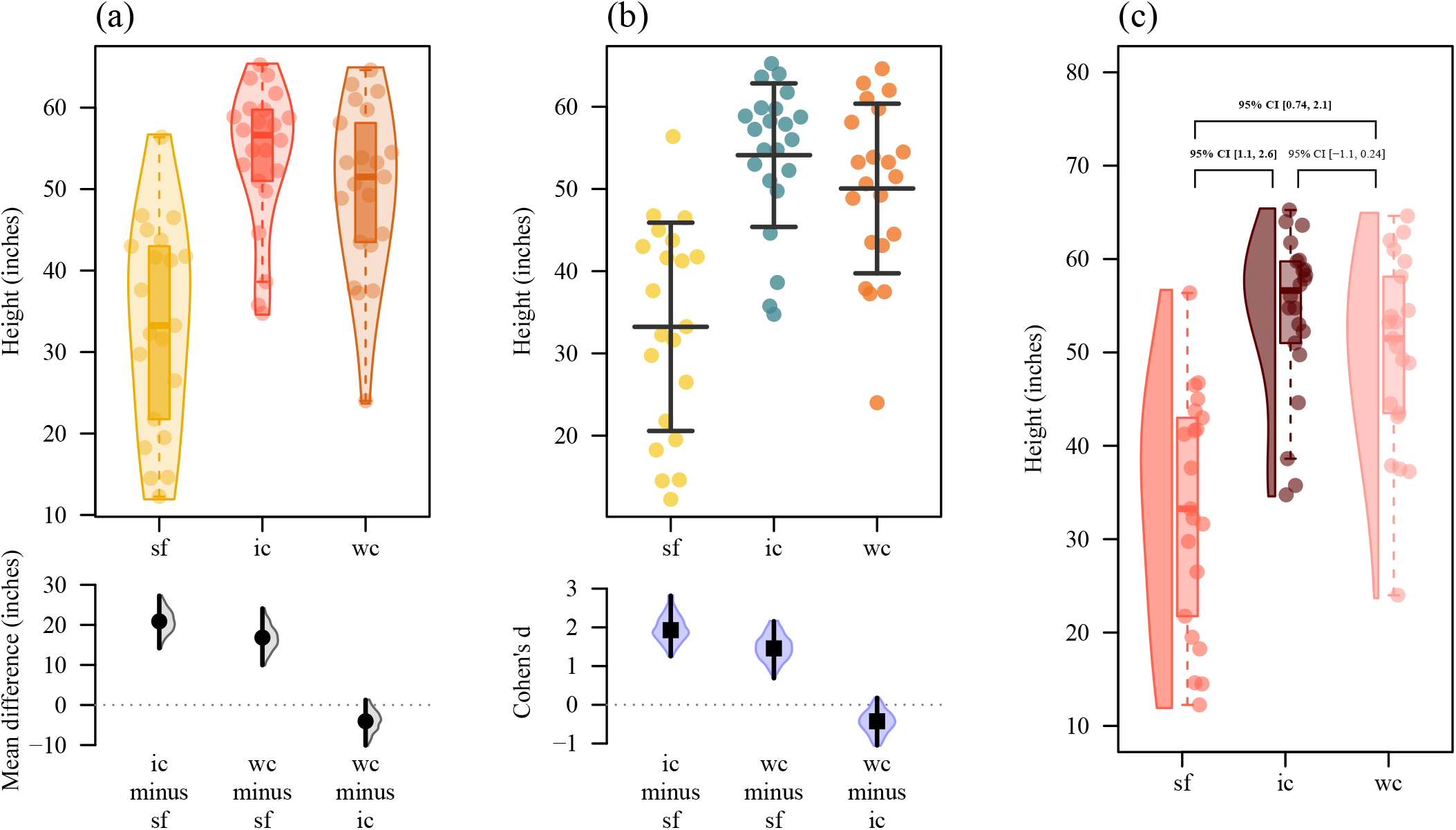
Cumming plot (A and B), and box-violin plot (C) showing height of self and cross-fertilised plants (sf = self-fertilised; ic = inter-crossed fertilised; wc = westerham-crossed fertilised). A) Top plot region represents group data and bottom region represents effect size statistics. Violins exhibit distribution of the data. The boxplots display the group median and the 75th and 25th percentiles. The whiskers extend to the minimum and maximum values, but exclude outliers that are beyond 1.5 times the interquartile range. Circles indicate individual values. Half violins in the lower region exhibit the distribution of bootstrapped differences, solid circles show mean difference and vertical bars show 95% confidence intervals of mean difference. B) Upper horizontal and vertical bars and points exhibit group data, and lower violins exhibit effect size statistics. The middle horizontal lines in the group data represent means, and vertical lines represent standard deviation. Circles indicate individual values. Violins on the lower region show the distribution of bootstrapped Cohen’s d (standardised mean difference), squares show Cohen’s d, and vertical bars show 95% confidence interval of Cohen’s d. C) Violins show distribution of the data in each group. The boxplots display median and the 75th and 25th percentiles. The whiskers extend to the minimum and maximum values, but exclude outliers that are beyond 1.5 times the interquartile range. Circles indicate individual values. Brackets show 95% confidence intervals of Hedges’ g for pairwise comparison.

Researchers could report the results by using the DurgaDiff output as: The height of the inter-crossed plants (n = 22, M = 54.11, SD = 8.73) was greater than the self-fertilised plants (n = 21, M = 33.23, SD = 12.66) (mean difference: 20.88, 95% CI [14.41, 27.69], Figure 3a); or (Cohen’s d = 1.92, 95% CI [1.10, 2.61], Figure 3b); or (Hedges’ g = 1.89, 95% CI = [1.15, 2.55], Figure 3c). Similarly, Westerham-crossed plants (n = 21, M = 50.05, SD = 10.31) were taller than self-fertilised plants (n = 21, M = 33.23, SD = 12.66) (mean difference: 16.82, 95% CI [10.01, 23.37], Figure 3a); or (Cohen’s d = 1.45, 95% CI [0.72, 2.13], Figure 3b); or (Hedges’ g = 1.43, 95% CI = [0.74, 2.08], Figure 3c). However, there was no evidence that the heights of inter-crossed plants (n = 22, M = 54.11, SD = 8.73) and Westerham-crossed plants (n = 21, M = 50.05, SD = 10.31) differed (mean difference: -4.05, 95% CI = [-10.01, 1.40], Figure 3a); or (Cohen’s d = -0.42, 95% CI [-1.0, 0.2], Figure 3b); or (Hedges’ g = - 0.42, 95% CI = [-1.01, 0.23], Figure 3c).

Examples of effect size calculation and visualisation of more than three groups are available via the package vignette.

## Conclusions

The Durga R package offers an easy way to estimate and plot effect sizes. The main novelty of the package is the ability to plot effect sizes together with a wide range of options for displaying group data. The strength of the package is its flexibility combined with an easy-to-use interface, which provides a creative platform for users to produce informative and aesthetic plots.

Durga performs post-hoc pairwise analysis and plotting for multi-group data sets. It does not determine whether there is evidence that measured values differ among groups. Researchers might perform ANOVA-like analyses, such as Anova, GLM or GLMM, and then use the R package `effectsize` (Ben-Shachar et al., 2020), or similar packages, to determine if there is evidence of differences among groups.

We hope that Durga can facilitate and encourage the uptake of estimation statistics and assist researchers in moving away from *p*-value driven dichotomous decision making in their research. We plan to add new functionality to this package to plot a wider range of data types, and to improve plot aesthetics. We encourage users to suggest new features and welcome users’ support to identify and fix issues.

## Acknowledgements

We acknowledge the *Wallumattagal clan of the Dharug nation* as the traditional custodians of the Macquarie University land. We thank Prof. Dr. Marie E. Herberstein for her support and mentorship. We thank our families for their continuous support and inspiration.

## Authors contributions

MKK conceived the idea, MKK and DJM designed the R package, DJM developed the software and unit tests, MKK collected and analysed data, MKK prepared the first draft of the manuscript, both authors contributed to developing the manuscript and gave final approval for publication.

## Data availability statement

All data used in the manuscript are installed with the Durga package. The Durga source code and data are available GitHub (https://github.com/KhanKawsar/EstimationPlot). R code used to generate the figures is available in the supplementary file S2 and via GitHub (https://github.com/JimMcL/Durga-paper/blob/main/R/manuscript_plot.R).

## Statement of Diversity and Inclusion

We strongly support equity, diversity and inclusion in science. The authors come from Bangladesh and Australia, and are early career researchers. One or more authors are from underrepresented ethnic minorities in science. We acknowledge the lack of gender diversity in the current project and are open for future collaboration to improve gender diversity in future projects.

## Notes

### Competing Interest Statement

The authors have declared no competing interest.

