## Supplementary figures and images for "Durga: An R package for effect size estimation and visualisation"

### Supporting Information S1

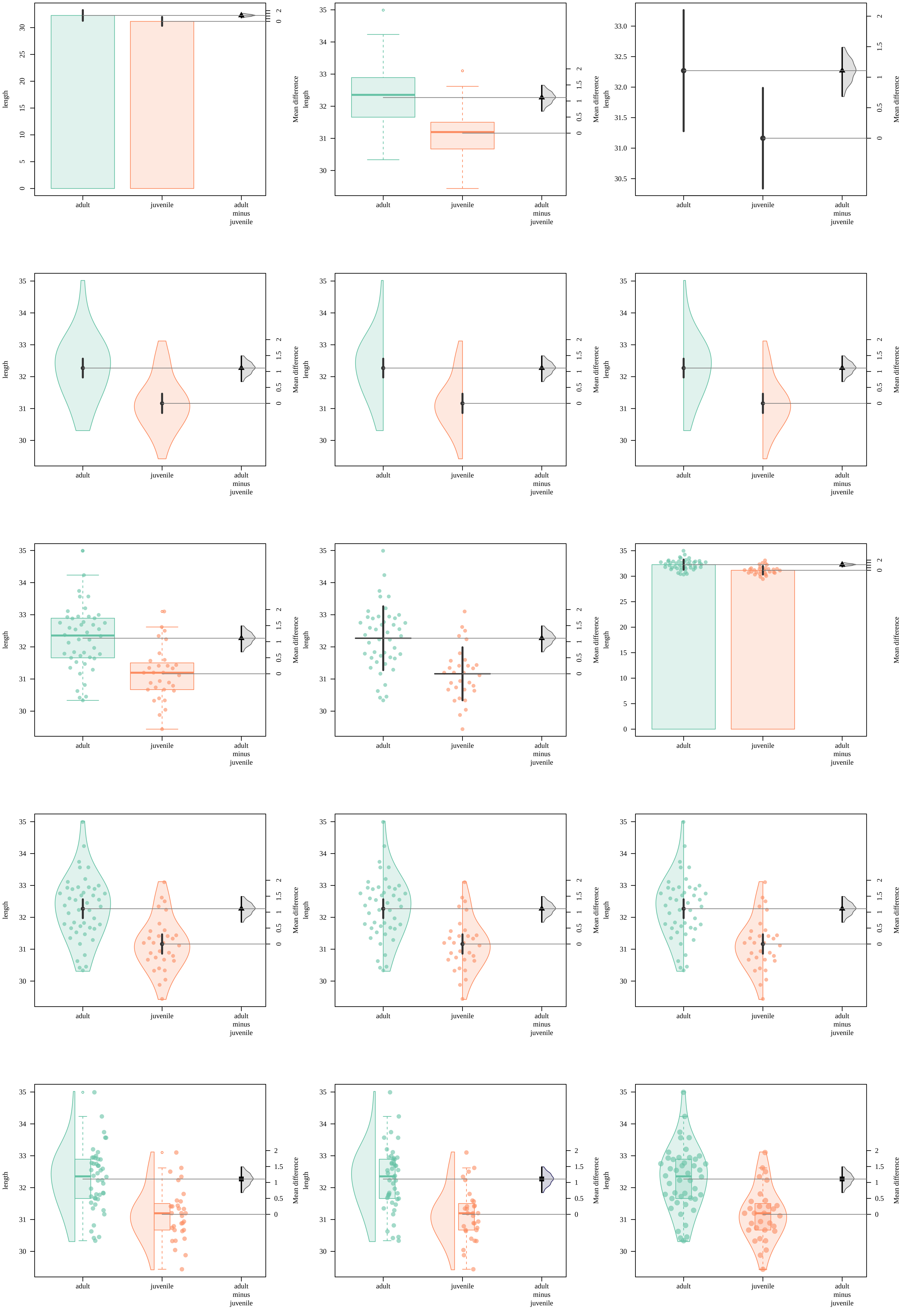

### Supporting Information S3

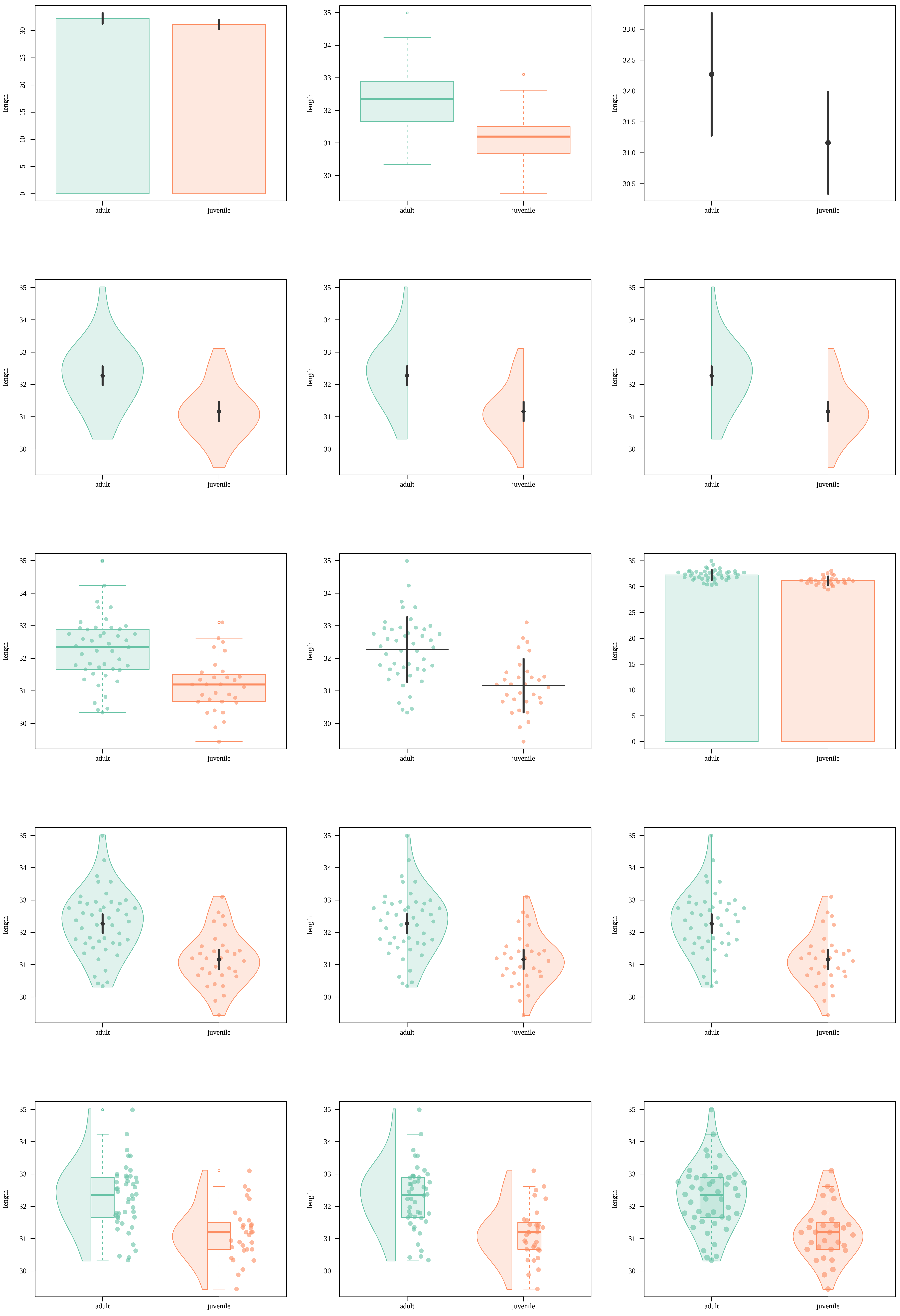

### Supporting Information S4

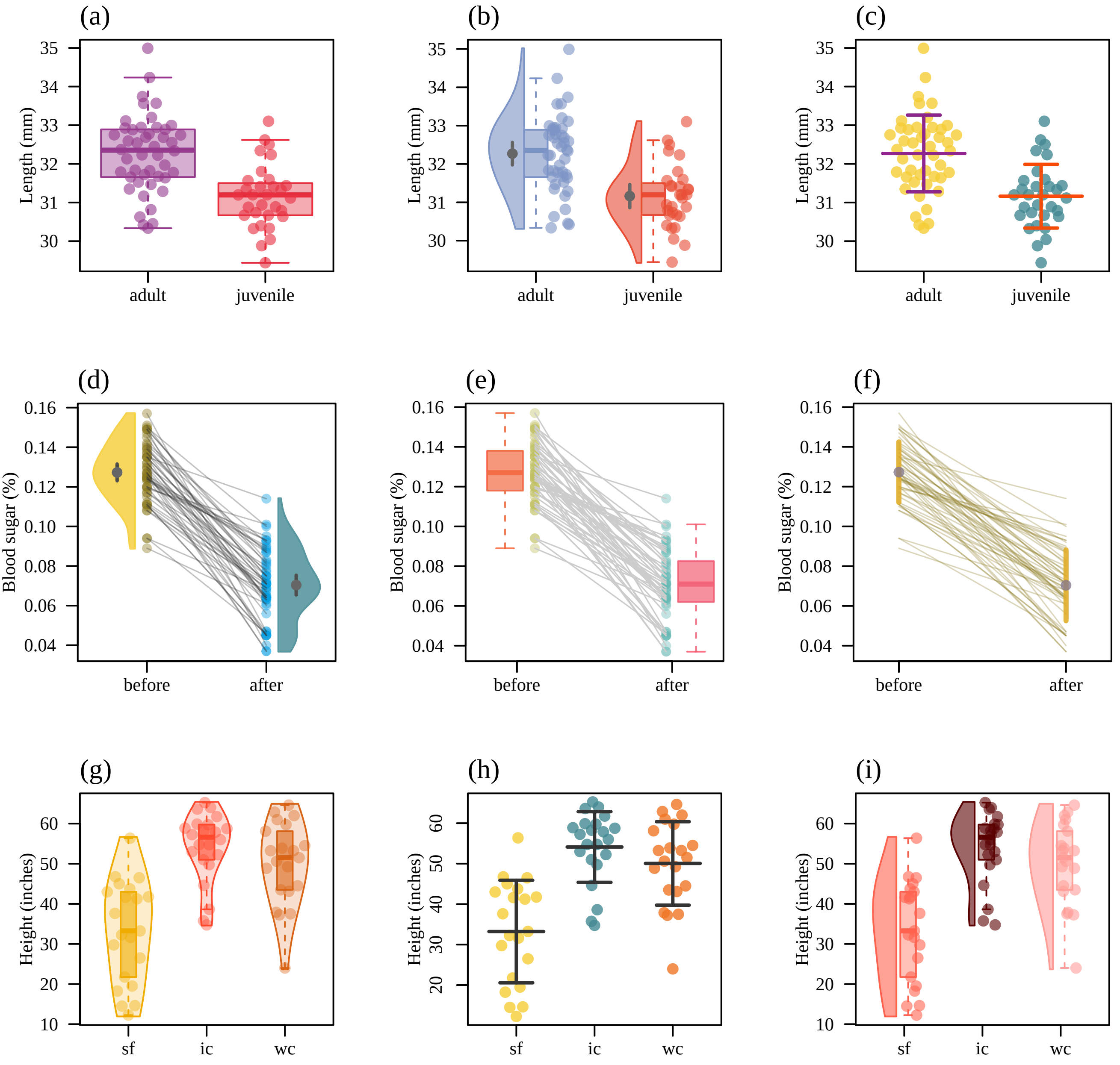
