## Supporting Information S2 for "Durga: An R package for effect size estimation and visualisation": manuscript_plot.R

```
# Code to generate the figures in the paper

#load package
#install.packages('Durga')

# to install the development version
#devtools::install_github("KhanKawsar/EstimationPlot", build_vignettes = TRUE)

## load library

library(Durga)


#### Colour palette for the manuscript plot ####

# to plot figure we used our own colour palette

## Based on tent inn the love umbrella
a <- rgb(240, 172, 1, max = 255) #
b <- rgb(254, 74, 47, max = 255)
c <- rgb(219, 101, 23, max = 255) #
d <- rgb(97, 79, 131, max = 255)
e <- rgb(113, 112, 114, max = 255)
f <- rgb(154, 92, 4, max = 255)
g <- rgb(180, 130, 216, max = 255)
h <- rgb(234, 44, 62, max = 255)

Louis <- c(a,b,c,d,e,f,g,h)

## based on house in love umbrella
a1 <- rgb(179, 162, 190, max = 255) #
b1 <- rgb(223, 173, 39, max = 255)
c1 <- rgb(220, 103, 110, max = 255)
d1 <- rgb(216, 86, 40, max = 255) #
e1 <- rgb(135, 122, 140, max = 255)
f1 <- rgb(254, 226, 166, max = 255)
g1 <- rgb(78, 73, 78, max = 255)
h1 <- rgb(220, 103, 110, max = 255)
i1 <- rgb(234, 44, 62, max = 255)

Home <- c(a1,b1,c1,d1,e1,f1,g1,h1, i1)

## Based on tractor
a2 <- rgb(124, 148, 198, max = 255) #
b2 <- rgb(233, 76, 51, max = 255)
c2 <- rgb(156, 206, 169, max = 255)
d2 <- rgb(245, 206, 53, max = 255) #
e2 <- rgb(68, 138, 148, max = 255)
g2 <- rgb(240, 118, 37, max = 255)
i2 <- rgb(252, 215, 138, max = 255)
f2 <- rgb(48, 45, 100, max = 255)
h2 <- rgb(186, 123, 82, max = 255)

Khona <- c(a2,b2,c2,d2,e2, g2,i2,f2,h2)

## Baes on tomorrow is a brand new day
a3 <- rgb(220, 212, 0, max = 255) #
b3 <- rgb(175, 160, 212, max = 255)
c3 <- rgb(156, 206, 169, max = 255)
d3 <- rgb(177, 152, 140, max = 255) #
e3 <- rgb(238, 191, 5, max = 255)
f3 <- rgb(78, 78, 78, max = 255)
g3 <- rgb(80, 131, 181, max = 255)
h3 <- rgb(222, 41, 169, max = 255)
i3 <- rgb(205, 191, 160, max = 255)
j3 <- rgb(222, 190, 221,	max = 255)
k3 <- rgb(220, 189, 159, max = 255)
m3 <- rgb(212,	64,	33,	max = 255)
n3 <- rgb(60,	165,	58, max = 255)
o3 <- rgb(232, 130, 41, max = 255)
p3 <- rgb(1,	135,	131,	max = 255)
q3 <- rgb(11,	179,	177, max = 255)
r3 <- rgb(193,	112,	46, max = 255)


Leelaboti <- c(a3, b3, c3, d3, e3, f3, g3,h3, i3, j3, k3, m3, n3, o3, p3, q3, r3)


## Britta teckentrup : big and small

a4 <- rgb(0,	111,	163	, max = 255) #
b4 <- rgb(231,	81,	44, max = 255)
c4 <- rgb(156, 206, 169, max = 255)
d4 <- rgb(158,	191,	72, max = 255) #
e4 <- rgb(149,	57,	140, max = 255)
f4 <- rgb(231,	46,	63, max = 255)
g4 <- rgb(228,	132,	156, max = 255)
h4 <- rgb(242,	104,	42, max = 255)

Britta <- c(a4, b4, c4, d4, e4, f4, g4, h4)

## Britta teckentrup : Blue

a5 <- rgb(247,	217,	8	, max = 255) #
b5 <- rgb(139,	121,	29, max = 255)
c5 <- rgb(3,	156,	225, max = 255)
d5 <- rgb(17,	79,	111, max = 255) #
e5 <- rgb(93,	111,	159, max = 255)
f5 <- rgb(6,	156,	119, max = 255)
g5 <- rgb(51,	51,	54, max = 255)


Blue <- c(a5, b5, c5, d5, e5, f5, g5)


## Britta teckentrup : Memory
a6 <- rgb(142,	101,	74, max = 255) #
b6 <- rgb(247,	76,	7, max = 255)
c6 <- rgb(240,	15,	12, max = 255)
d6 <- rgb(107,	111,	22, max = 255) #
e6 <- rgb(209,	94,	8, max = 255)
f6 <- rgb(14,	103,	108, max = 255)
g6 <- rgb(183,	96,	28, max = 255)


Memory <- c(a6, b6, c6, d6, e6, f6, g6)


## Britta teckentrup :Before i woke up

a7 <- rgb(117,	106,	55, max = 255) #
b7 <- rgb(55,	48,	48, max = 255)
c7 <- rgb(53,	68,	47, max = 255)
d7 <- rgb(96,	90,	72, max = 255) #
e7 <- rgb(112,	123,	128, max = 255)
f7 <- rgb(74,	78,	52, max = 255)
g7 <- rgb(192,	152,	97, max = 255)

Dream <- c(a7, b7, c7, d7, e7, f7, g7)

## Boishakh: from Horse

a8 <- rgb(222,	49,	34, max = 255) #
b8 <- rgb(255,	99,	79, max = 255)
c8 <- rgb(90,	1,	2, max = 255)
d8 <- rgb(255,	158,	152, max = 255) #
e8 <- rgb(248,	171,	44, max = 255)
f8 <- rgb(145,	208,	253, max = 255)
g8 <- rgb(3,	65,	73, max = 255)
h8 <- rgb(189,	106,	253, max = 255)

Boishakh <- c(a8, b8, c8, d8, e8, f8, g8, h8)

## Boishakh from bird

a9 <- rgb(243,	106,	65, max = 255) #
b9 <- rgb(243,	96,	118, max = 255)
c9 <- rgb(253,	215,	87, max = 255)
d9 <- rgb(242,	133,	90, max = 255) #
e9 <- rgb(192,	192,	97, max = 255)
f9 <- rgb(103,	186,	181, max = 255)
g9 <- rgb(191,	228,	213, max = 255)
h9 <- rgb(90,	165,	78, max = 255)

Boishakh2 <- c(a9, b9, c9, d9, e9, f9, g9, h9)

## Boishakh from mask


a10 <- rgb(39,	34,	90, max = 255) #
b10 <- rgb(100,	77,	159, max = 255)
c10 <- rgb(237,	61,	55, max = 255)
d10 <- rgb(197,	88,	163, max = 255) #
e10 <- rgb(241,	103, 56, max = 255)
f10 <- rgb(250,	235,	75, max = 255)


Boishakh3 <- c(a10, b10, c10, d10, e10, f10)


## coco: shades of grey

Coco <- c("grey20", "grey20", "grey40", "grey50", "grey60", "grey70", "grey80", "grey90")

## Pride: colours of rainbow flag

a11 <- rgb(141,	39,	141, max = 255) #
b11 <- rgb(4,	77,	163, max = 255)
c11 <- rgb(85,	185,	72, max = 255)
d11 <- rgb(255,	216,	0, max = 255) #
e11 <- rgb(192,	192,	97, max = 255)
f11 <- rgb(238,	26,	37, max = 255)
g11 <- rgb(254,	225,	238, max = 255)
h11 <- rgb(245,	154,	193, max = 255)
i11 <- rgb(156,	207,	240, max = 255)
j11 <- rgb(99,	64,	46, max = 255)
k11 <- rgb(0,	0,	0, max = 255)


#### all palletes

Leelaboti <- c(a3, b3, c3, d3, e3, f3, g3,h3, i3, j3, k3, m3, n3, o3, p3, q3, r3)
Khona <- c(a2, b2, c2, d2, e2, g2,i2,f2,h2)
Louis <- c(a,b,c,d,e,f,g,h)
Britta <- c(a4, b4, c4, d4, e4, f4, g4, h4)
Coco <- c("grey20", "grey20", "grey40", "grey50", "grey60", "grey70", "grey80", "grey90")
Home <- c(a1,b1,c1,d1,e1,f1,g1,h1, i1)
Blue <- c(a5, b5, c5, d5, e5, f5, g5)
Memory <- c(a6, b6, c6, d6, e6, f6, g6)
Dream <- c(a7, b7, c7, d7, e7, f7, g7)
Boishakh <- c(a8, b8, c8, d8, e8, f8, g8, h8)
Boishakh2 <- c(a9, b9, c9, d9, e9, f9, g9, h9)
Boishakh3 <- c(a10, b10, c10, d10, e10, f10)
Pride <- c(a11, b11, c11, d11, e11, f11, g11, h11, i11, j11, k11)


#### manuscript plot code ####

## load data
data(damselfly)

# calculate effect size

d1 <- DurgaDiff(damselfly, data.col = 1, group.col = 3, effect.type = "mean",
                contrasts = "adult - juvenile")
d2 <- DurgaDiff(damselfly, data.col = 1, group.col = 3, effect.type = "cohens",
                contrasts = "adult - juvenile")
d3 <- DurgaDiff(damselfly, data.col = 1, group.col = 3, effect.type = "hedges",
                contrasts = "adult - juvenile")


#### Figure 1 ####
pageWidthLarge<- 7.08661
pageHeightLarge <- pageWidthLarge * 0.45
pagePaper <- 'special'
fontFamily <- 'Times'

# Create output folder if it doesn't already exist
if (!dir.exists("../output"))
  dir.create("../output")

png("../output/Figure 1.png", width=pageWidthLarge,
    height= pageHeightLarge, family=fontFamily,
    units= "in",type= "cairo",res = 2400)

#To download pdf follow the code
#pdf("../output/Figure 1.pdf", width=pageWidthLarge,
 #   height= pageHeightLarge, family=fontFamily, paper=pagePaper)


layout(matrix(c(1,2,3), nrow=1,ncol=3,byrow =  TRUE))
par(mgp = c(2,1,0))


##
p1 <- DurgaPlot(d1, box = DurgaTransparent(Britta[5:6],0.0),
                box.fill = DurgaTransparent(Britta[5:6],0.6),
                points = DurgaTransparent(Britta[5:6],0.4),
                points.params = list(cex = 1.2),
                ef.size.pch = 15,
                ef.size = DurgaTransparent(Khona[6], 0.0),
                ef.size.violin = DurgaTransparent(Khona[6], 0.4),
                ef.size.mean.line.dx = 0.4,
                las = 1,
                ef.size.params = list(las = 1),
                box.outline = F,
                left.ylab = "Length (mm)",
                ef.size.label = "Mean difference (mm)")

mtext("(a)", side=3, line=0.5, adj=0, cex=1.0)

p2 <- DurgaPlot(d2, box = DurgaTransparent(Khona,0.0),
                box.fill = DurgaTransparent(Khona,0.4),
                box.outline = FALSE,
                box.params = list(boxwex = 0.2),
                violin = DurgaTransparent(Khona,0),
                violin.fill = DurgaTransparent(Khona,0.4),
                violin.width = 0.30,
                violin.dx = -0.1,
                #box.dx = 0.25,
                points = DurgaTransparent(Khona,0.4),
                points.dx = 0.20,
                points.method = "jitter",
                points.params = list(cex = 1.2),
                las = 1,
                ef.size.params = list(las = 1),
                #ef.size.dx = 0.6,
                ef.size.pch = 16,
                ef.size.violin = DurgaTransparent(Leelaboti[2], 0.0),
                #ef.size = DurgaTransparent(Leelaboti[12], 0.0),
                ef.size = TRUE,
                ef.size.violin.shape = "full",
                ef.size.mean.line.dx = -0.18,
                central.tendency = "grey40",
                central.tendency.type = "mean",
                central.tendency.symbol = "point",
                central.tendency.dx = -0.20,
                error.bars.lwd = 2,
                error.bars = "grey40",
                error.bars.type = "CI",
                left.ylab = "Length (mm)")

mtext("(b)", side=3, line=0.5, adj=0, cex=1.0)

DurgaPlot(d3,
          box = FALSE,
          violin = FALSE,
          points = DurgaTransparent(Khona[4:5],0.2),
          central.tendency = DurgaTransparent(c(Pride[1],Memory[2]),0.0),
          points.params = list(cex = 1.2),
          central.tendency.type = "mean",
          central.tendency.symbol = "segment",
          central.tendency.params = list(lwd = 2.0), #does not seem to work
          error.bars = DurgaTransparent(c(Pride[1],Memory[2]),0.0),
          error.bars.type = "SD",
          error.bars.cross.width = 0.1,
          error.bars.lwd = 2,
          ef.size.violin = DurgaTransparent("blue", 0.6),
          ef.size = TRUE,
          ef.size.mean.line.dx = 0.35,
          las = 1,
          ef.size.params = list(las = 1),
          left.ylab = "Length (mm)")

mtext("(c)", side=3, line=0.5, adj=0, cex=1.0)

dev.off()

#### Figure 2 ####

# insulin data analysis
data(insulin)
head(insulin)

d4 <- DurgaDiff(insulin, data.col = 1, group.col = 2, id.col = 3, effect.type = "mean", groups = c("before", "after"))
d5 <- DurgaDiff(insulin, data.col = 1, group.col = 2, id.col = 3, effect.type = "cohens", groups = c("before", "after"))
d6 <- DurgaDiff(insulin, data.col = 1, group.col = 2, id.col = 3, effect.type = "hedges", groups = c("before", "after"))


pageWidthLarge<- 7.08661
pageHeightLarge <- pageWidthLarge * 0.45
pagePaper <- 'special'
fontFamily <- 'Times'


png("../output/Figure 2.png", width=pageWidthLarge,
    height= pageHeightLarge, family=fontFamily,
    units= "in",type= "cairo",res = 2400)

#To download pdf run the code
#pdf("../output/Figure 2.pdf", width = pageWidthLarge,
 #   height = pageHeightLarge, family = fontFamily, paper = pagePaper)


layout(matrix(c(1,2,3), nrow=1,ncol=3,byrow =  TRUE))

op <- par(mar = c(5, 4, 4, 5))
DurgaPlot(d4,
          violin = DurgaTransparent(Khona[4:5],0.1),
          violin.shape = c("left-half", "right-half"),
          violin.fill = DurgaTransparent(Khona[4:5],0.2),
          violin.adj = 1.2,
          violin.dx = c(-0.1,0.1),
          points = DurgaTransparent(Blue[2:3],0.6),
          las = 1,
          ef.size.params = list(las = 1, mgp = c(3.5, 1, 0)),
          ef.size.violin = DurgaTransparent(Louis,0.4),
          ef.size = Louis[3],
          ef.size.mean.line.dx = c(-0.21, 0.29),
          paired.lwd = 0.8,
          central.tendency = "grey40",
          central.tendency.dx = c(-0.25, 0.25),
          error.bars = "grey30",
          error.bars.lwd = 2,
          left.ylab = "Blood sugar (%)",
          ef.size.label = "Mean difference (%)")

mtext("(a)", side=3, line=0.5, adj=0, cex=1.0)
par(op)


DurgaPlot(d5,
          box = DurgaTransparent(Boishakh2, 0.1),
          box.fill = DurgaTransparent(Boishakh2, 0.3),
          box.dx = c(-0.25, 0.35),
          box.outline = FALSE,
          box.params = list(boxwex = 0.3),
          points = DurgaTransparent(Boishakh2[5:6], 0.6),
          points.dx = c(-0.0,0.10),
          x.axis.dx = c(-0.15,0.15),
          paired = "grey80",
          paired.lwd = 0.8,
          ef.size.pch = 18,
          ef.size.params = list(las = 1),
          ef.size = DurgaTransparent(Blue[4], 0.1),
          ef.size.violin = DurgaTransparent(Blue[3], 0.7),
          ef.size.violin.shape = "full",
          ef.size.mean.line.dx = c(-0.10, 0.50),
          error.bars.lwd = 2,
          las = 1,
          left.ylab = "Blood sugar (%)")

mtext("(b)", side=3, line=0.5, adj=0, cex=1.0)

DurgaPlot(d6,
          box = FALSE,
          violin = FALSE,
          group.dx = c(-0.2,0.2),
          central.tendency = DurgaTransparent(Home[5], 0.2),
          points = FALSE,
          #points.dx = c(0.1,-0.1),
          error.bars = DurgaTransparent(Home[2], 0.1),
          error.bars.type = "SD",
          paired = DurgaTransparent(Blue[2], 0.7),
          paired.lwd = 0.8,
          ef.size = DurgaTransparent("purple", 0.2),
          ef.size.violin = DurgaTransparent("red", 0.8),
          ef.size.pch = 15,
          ef.size.params = list(las = 1),
          las =1,
          left.ylab = "Blood sugar (%)")

mtext("(c)", side=3, line=0.5, adj=0, cex=1.0)

dev.off()


#### Figure 3 ####

d7 <- DurgaDiff(petunia, data.col = 1, group.col = 2,
                groups = c("sf" = "self_fertilised", "ic" = "inter_cross", "wc" = "westerham_cross"))
d8 <- DurgaDiff(petunia, data.col = 1, group.col = 2,
                groups = c("sf" = "self_fertilised", "ic" = "inter_cross", "wc" = "westerham_cross"),
                effect.type = "cohens")
d9 <- DurgaDiff(petunia, data.col = 1, group.col = 2,
                groups = c("sf" = "self_fertilised", "ic" = "inter_cross", "wc" = "westerham_cross"),
                effect.type = "hedges")


###
pageWidthLarge<- 7.08661
pageHeightLarge <- pageWidthLarge * 0.65
pagePaper <- 'special'
fontFamily <- 'Times'


png("../output/Figure 3.png", width=pageWidthLarge,
    height= pageHeightLarge, family=fontFamily,
    units= "in",type= "cairo",res = 2400)

# to download pdf figure, run the code
#pdf("../output/Figure 3.pdf", width=pageWidthLarge,
   # height= pageHeightLarge, family=fontFamily, paper=pagePaper)


layout(matrix(c(1,2,3), nrow=1,ncol=3,byrow =  TRUE))

par(mgp = c(2.5,1,0), mar = c(6,4,4,2) + 0.1)

DurgaPlot(d7,
          contrasts = "ic - sf,  wc - sf, wc - ic",
          box = DurgaTransparent(Louis,0.0),
          box.fill = DurgaTransparent(Louis,0.4),
          box.outline = FALSE,
          box.params = list(boxwex = 0.2),
          violin.shape = "full",
          violin = DurgaTransparent(Louis,0),
          violin.fill = DurgaTransparent(Louis,0.8),
          violin.width = 0.30,
          points = DurgaTransparent(Louis,0.6),
          points.params = list(cex = 1.2),
          las = 1,
          ef.size.dx = c(-1.0, -1.0, 0.0),
          ef.size.params = list(las = 1, mgp = c(2.5,1,0)),
          ef.size.pch = 16,
          ef.size.height = 0.35,
          ef.size.ticks = c(-10,0,10,20,30),
          left.ylab = "Height (inches)",
          ef.size.label = "Mean difference (inches)")

mtext("(a)", side=3, line=0.5, adj=0, cex=1.0)

#b
par(mgp = c(2.5,1,0), mar = c(6,4,4,2) + 0.1)

DurgaPlot(d8, contrasts = "ic - sf,  wc - sf, wc - ic",
          box = FALSE,
          violin = FALSE,
          points = DurgaTransparent(Khona[4:6],0.2),
          points.params = list(cex = 1.2),
          central.tendency.type = "mean",
          central.tendency.symbol = "segment",
          central.tendency.params = list(cex =2.5), #does not seem to work
          error.bars.type = "SD",
          error.bars.cross.width = 0.1,
          error.bars.lwd = 2,
          ef.size.violin = DurgaTransparent("blue", 0.6),
          ef.size.violin.shape = "full",
          ef.size = TRUE,
          las = 1,
          ef.size.params = list(las = 1, mgp = c(2.5,1,0)),
          ef.size.dx = c(-1.0, -1.0, 0.0),
          ef.size.pch = 15,
          left.ylab = "Height (inches)")

mtext("(b)", side=3, line=0.5, adj=0, cex=1.0)

# c
#par(mgp = c(2,1,0), mar = c(5,4,6,2) + 0.1)
p <- DurgaPlot(d9, contrasts = "ic - sf,  wc - sf, wc - ic",
               box = DurgaTransparent(Boishakh[2:4],0.0),
               box.fill = DurgaTransparent(Boishakh[2:4],0.6),
               box.outline = FALSE,
               box.params = list(boxwex = 0.2),
               violin = DurgaTransparent(Boishakh[2:4],0),
               violin.fill = DurgaTransparent(Boishakh[2:4],0.4),
               violin.width = 0.30,
               violin.dx = -0.1,
               box.dx = 0.05,
               points = DurgaTransparent(Boishakh[2:4],0.4),
               points.dx = 0.10,
               points.method = "jitter",
               points.params = list(cex = 1.2),
               las = 1,
               ef.size.params = list(las = 1),
               ef.size = F,
               ef.size.pch = 15,
               ef.size.violin = DurgaTransparent(Boishakh[5], 0.0),
               ylim = c(10,80),
               left.ylab = "Height (inches)")

DurgaBrackets(p, lb.cex = 0.5, br.lwd = 0.8, br.col = "grey5", lb.col = "grey5")

mtext("(c)", side=3, line=0.5, adj=0, cex=1.0)

dev.off()


#### Supplementary figure S1 ####

pageWidthLarge<- 14.08661
pageHeightLarge <- pageWidthLarge * 1.5
pagePaper <- 'special'
fontFamily <- 'Times'


png("../output/Supporting Information S1.png", width=pageWidthLarge,
    height= pageHeightLarge, family=fontFamily,
    units= "in",type= "cairo",res = 1200)

# to download pdf figure, run the code
#pdf("../output/Supporting Information S1.pdf", width=pageWidthLarge,
    #height= pageHeightLarge, family=fontFamily, paper=pagePaper)


layout(matrix(c(1,2,3,4,5,6,7,8,9,10,11,12,13,14,15), nrow=5,ncol=3,byrow =  TRUE))

DurgaPlot(d1, bar = T, points = FALSE,
          error.bars.type = "SD")

DurgaPlot(d1, box = T,las = 1,
          points = FALSE)


DurgaPlot(d1, box = FALSE, violin = FALSE,
          central.tendency.type = "mean",
          central.tendency.symbol = "point",
          central.tendency = TRUE,
          error.bars.type = "SD",
          central.tendency.params = list(cex = 1.5),
          points = FALSE,
           las = 1)


DurgaPlot(d1, violin = TRUE, violin.shape = "full",
          points =FALSE,  las = 1)

DurgaPlot(d1, violin = TRUE,  points = FALSE, las = 1)

DurgaPlot(d1, violin = TRUE, violin.shape = "right-half",
          points =FALSE,  las = 1)

DurgaPlot(d1, box = TRUE,  las = 1)

DurgaPlot(d1, box = FALSE, violin = FALSE,
          central.tendency.type = "mean",
          central.tendency.symbol = "segment",
          central.tendency = TRUE,
          error.bars.type = "SD",
          las = 1)

DurgaPlot(d1, bar = TRUE,  error.bars.type = "SD",  las = 1)
DurgaPlot(d1, violin = TRUE, violin.shape = "full", las = 1)
DurgaPlot(d1, violin = TRUE, violin.shape = "right-half",  las = 1)
DurgaPlot(d1, violin = TRUE,  las =1 )

DurgaPlot(d1, box = T,
          box.params = list(boxwex = 0.2),
          violin = T,
          violin.width = 0.30,
          violin.dx = -0.1,
          points = T,
          points.dx = 0.20,
          points.method = "jitter",
          points.params = list(cex = 1.2),
          ef.size.pch = 15,
          ef.size.violin = T,
          las = 1,
          ef.size.params = list(las = 1))

DurgaPlot(d1, box = T,
          box.outline = FALSE,
          box.params = list(boxwex = 0.2),
          violin = T,
          violin.width = 0.30,
          violin.dx = -0.1,
          box.dx = 0.05,
          points = T,
          points.dx = 0.10,
          points.method = "jitter",
          points.params = list(cex = 1.2),
          ef.size.pch = 15,
          ef.size.violin = DurgaTransparent(Khona[8], 0.0),
          las = 1,
          ef.size.params = list(las = 1))


DurgaPlot(d1, box = T,
          box.outline = FALSE,
          box.params = list(boxwex = 0.2),
          violin.shape = "full",
          violin = T,
          violin.width = 0.30,
          points = T,
          points.params = list(cex = 1.5),
          ef.size.pch = 15,
          ef.size.violin = T,
          las = 1,
          ef.size.params = list(las = 1))

dev.off()


#### Supplementary figure S3 ####

pageWidthLarge<- 14.08661
pageHeightLarge <- pageWidthLarge * 1.5
pagePaper <- 'special'
fontFamily <- 'Times'

#to download pdf figure run this code
#pdf("../output/Supporting Information S3.pdf", width=pageWidthLarge,
 #   height= pageHeightLarge, family=fontFamily, paper=pagePaper)

png("../output/Supporting Information S3.png", width=pageWidthLarge,
    height= pageHeightLarge, family=fontFamily,
    units= "in",type= "cairo",res = 1200)

layout(matrix(c(1,2,3,4,5,6,7,8,9,10,11,12,13,14,15), nrow=5,ncol=3,byrow =  TRUE))

DurgaPlot(d1, bar = T, points = FALSE,
          error.bars.type = "SD", ef.size = FALSE)

DurgaPlot(d1, box = T,las = 1,
          points = FALSE, ef.size = FALSE)


DurgaPlot(d1, box = FALSE, violin = FALSE,
          central.tendency.type = "mean",
          central.tendency.symbol = "point",
          central.tendency = TRUE,
          error.bars.type = "SD",
          central.tendency.params = list(cex = 1.5),
          points = FALSE,
          las = 1,
          ef.size = FALSE)


DurgaPlot(d1, violin = TRUE, violin.shape = "full",
          points =FALSE,  las = 1, ef.size = FALSE)

DurgaPlot(d1, violin = TRUE, points = FALSE, las = 1, ef.size = FALSE)

DurgaPlot(d1, violin = TRUE, violin.shape = "right-half",
          points =FALSE, las = 1, ef.size = FALSE, )

DurgaPlot(d1, box = TRUE, las = 1, ef.size = FALSE)

DurgaPlot(d1, box = FALSE, violin = FALSE,
          central.tendency.type = "mean",
          central.tendency.symbol = "segment",
          central.tendency = TRUE,
          error.bars.type = "SD",
          las = 1, ef.size = FALSE)

DurgaPlot(d1, bar = TRUE,  error.bars.type = "SD",las = 1,
          ef.size = FALSE,)
DurgaPlot(d1, violin = TRUE, violin.shape = "full", las = 1,
          ef.size = FALSE,)
DurgaPlot(d1, violin = TRUE, violin.shape = "right-half",  las = 1,
          ef.size = FALSE)
DurgaPlot(d1, violin = TRUE, las =1,
          ef.size = FALSE)

DurgaPlot(d1, box = T,
          box.params = list(boxwex = 0.2),
          violin = T,
          violin.width = 0.30,
          violin.dx = -0.1,
          points = T,
          points.dx = 0.20,
          points.method = "jitter",
          points.params = list(cex = 1.2),
          las = 1,
          ef.size = FALSE)

DurgaPlot(d1, box = T,
          box.outline = FALSE,
          box.params = list(boxwex = 0.2),
          violin = T,
          violin.width = 0.30,
          violin.dx = -0.1,
          box.dx = 0.05,
          points = T,
          points.dx = 0.10,
          points.method = "jitter",
          points.params = list(cex = 1.2),
          las = 1,
          ef.size = FALSE)


DurgaPlot(d1, box = T,
          box.outline = FALSE,
          box.params = list(boxwex = 0.2),
          violin.shape = "full",
          violin = T,
          violin.width = 0.30,
          points = T,
          points.params = list(cex = 1.5),
          las = 1,
          ef.size = FALSE)

dev.off()


#### Supplementary Figure S4 ####


pageWidthLarge <- 7.08661
pageHeightLarge <- pageWidthLarge * 1.0
pagePaper <- 'special'
fontFamily <- 'Times'


png("../output/Supporting Information S4.png", width=pageWidthLarge,
    height= pageHeightLarge, family=fontFamily,
    units= "in",type= "cairo",res = 2400)

#to downlaod pdf figure run this code
#pdf("../output/Supporting Information S4.pdf", width=pageWidthLarge,
#   height= pageHeightLarge, family=fontFamily, paper=pagePaper)


layout(matrix(c(1,2,3,4,5,6,7,8,9), nrow=3,ncol=3,byrow =  TRUE))
par(mgp = c(2,1,0), mar = c(c(4, 4, 3, 2) + 0.1))
##
p1 <- DurgaPlot(d1, box = DurgaTransparent(Britta[5:6],0.0),
                box.fill = DurgaTransparent(Britta[5:6],0.6),
                points = DurgaTransparent(Britta[5:6],0.4),
                points.params = list(cex = 1.2),
                ef.size = F,
                las = 1,
                box.outline = F,
                left.ylab = "Length (mm)",
)
mtext("(a)", side=3, line=0.5, adj=0, cex=1.0)

p2 <- DurgaPlot(d2, box = DurgaTransparent(Khona,0.0),
                box.fill = DurgaTransparent(Khona,0.4),
                box.outline = FALSE,
                box.params = list(boxwex = 0.2),
                violin = DurgaTransparent(Khona,0),
                violin.fill = DurgaTransparent(Khona,0.4),
                violin.width = 0.30,
                violin.dx = -0.1,
                #box.dx = 0.25,
                points = DurgaTransparent(Khona,0.4),
                points.dx = 0.20,
                points.method = "jitter",
                points.params = list(cex = 1.2),
                las = 1,
                ef.size = FALSE,
                central.tendency = "grey40",
                central.tendency.type = "mean",
                central.tendency.symbol = "point",
                central.tendency.dx = -0.20,
                error.bars.lwd = 2,
                error.bars = "grey40",
                error.bars.type = "CI",
                left.ylab = "Length (mm)")

mtext("(b)", side=3, line=0.5, adj=0, cex=1.0)

DurgaPlot(d3,
          box = FALSE,
          violin = FALSE,
          points = DurgaTransparent(Khona[4:5],0.2),
          central.tendency = DurgaTransparent(c(Pride[1],Memory[2]),0.0),
          points.params = list(cex = 1.2),
          central.tendency.type = "mean",
          central.tendency.symbol = "segment",
          central.tendency.params = list(lwd = 2.0), #does not seem to work
          error.bars = DurgaTransparent(c(Pride[1],Memory[2]),0.0),
          error.bars.type = "SD",
          error.bars.cross.width = 0.1,
          error.bars.lwd = 2,
          ef.size = FALSE,
          las = 1,
          left.ylab = "Length (mm)")

mtext("(c)", side=3, line=0.5, adj=0, cex=1.0)

par(mgp = c(2.7,1,0),mar = c(4, 4, 2, 2))

DurgaPlot(d4,
          violin = DurgaTransparent(Khona[4:5],0.1),
          violin.shape = c("left-half", "right-half"),
          violin.fill = DurgaTransparent(Khona[4:5],0.2),
          violin.adj = 1.2,
          violin.dx = c(-0.1,0.1),
          points = DurgaTransparent(Blue[2:3],0.6),
          las = 1,
          ef.size = FALSE,
          paired.lwd = 0.8,
          central.tendency = "grey40",
          central.tendency.dx = c(-0.25, 0.25),
          error.bars = "grey30",
          error.bars.lwd = 2,
          left.ylab = "Blood sugar (%)")

mtext("(d)", side=3, line=0.5, adj=0, cex=1.0)

par(mgp = c(2.7,1,0),mar = c(4, 4, 2, 2))

DurgaPlot(d5,
          box = DurgaTransparent(Boishakh2, 0.1),
          box.fill = DurgaTransparent(Boishakh2, 0.3),
          box.dx = c(-0.25, 0.35),
          box.outline = FALSE,
          box.params = list(boxwex = 0.3),
          points = DurgaTransparent(Boishakh2[5:6], 0.6),
          points.dx = c(-0.0,0.10),
          x.axis.dx = c(-0.15,0.15),
          paired = "grey80",
          paired.lwd = 0.8,
          ef.size = FALSE,
          error.bars.lwd = 2,
          las = 1,
          left.ylab = "Blood sugar (%)")

mtext("(e)", side=3, line=0.5, adj=0, cex=1.0)

par(mgp = c(2.7,1,0),mar = c(4, 4, 2, 2))
DurgaPlot(d6,
          box = FALSE,
          violin = FALSE,
          group.dx = c(-0.2,0.2),
          central.tendency = DurgaTransparent(Home[5], 0.2),
          points = FALSE,
          #points.dx = c(0.1,-0.1),
          error.bars = DurgaTransparent(Home[2], 0.1),
          error.bars.type = "SD",
          paired = DurgaTransparent(Blue[2], 0.7),
          paired.lwd = 0.8,
          ef.size = FALSE,
          las =1,
          left.ylab = "Blood sugar (%)")

mtext("(f)", side=3, line=0.5, adj=0, cex=1.0)

par(mgp = c(2,1,0), mar = c(5,4,2,2) + 0.1)

DurgaPlot(d7,
          contrasts = "ic - sf,  wc - sf, wc - ic",
          box = DurgaTransparent(Louis,0.0),
          box.fill = DurgaTransparent(Louis,0.4),
          box.outline = FALSE,
          box.params = list(boxwex = 0.2),
          violin.shape = "full",
          violin = DurgaTransparent(Louis,0),
          violin.fill = DurgaTransparent(Louis,0.8),
          violin.width = 0.30,
          points = DurgaTransparent(Louis,0.6),
          points.params = list(cex = 1.2),
          las = 1,
          ef.size = FALSE,
          left.ylab = "Height (inches)")

mtext("(g)", side=3, line=0.5, adj=0, cex=1.0)

#b

DurgaPlot(d8, contrasts = "ic - sf,  wc - sf, wc - ic",
          box = FALSE,
          violin = FALSE,
          points = DurgaTransparent(Khona[4:6],0.2),
          points.params = list(cex = 1.2),
          central.tendency.type = "mean",
          central.tendency.symbol = "segment",
          central.tendency.params = list(cex =2.5), #does not seem to work
          error.bars.type = "SD",
          error.bars.cross.width = 0.1,
          error.bars.lwd = 2,
          ef.size = FALSE,
          left.ylab = "Height (inches)")

mtext("(h)", side=3, line=0.5, adj=0, cex=1.0)

# c
#par(mgp = c(2,1,0), mar = c(5,4,6,2) + 0.1)
p <- DurgaPlot(d9, contrasts = "ic - sf,  wc - sf, wc - ic",
               box = DurgaTransparent(Boishakh[2:4],0.0),
               box.fill = DurgaTransparent(Boishakh[2:4],0.6),
               box.outline = FALSE,
               box.params = list(boxwex = 0.2),
               violin = DurgaTransparent(Boishakh[2:4],0),
               violin.fill = DurgaTransparent(Boishakh[2:4],0.4),
               violin.width = 0.30,
               violin.dx = -0.1,
               box.dx = 0.05,
               points = DurgaTransparent(Boishakh[2:4],0.4),
               points.dx = 0.10,
               points.method = "jitter",
               points.params = list(cex = 1.2),
               las = 1,
               ef.size = F,
               left.ylab = "Height (inches)")


mtext("(i)", side=3, line=0.5, adj=0, cex=1.0)

dev.off()
```
